# The nuclear receptor RORα preserves cardiomyocyte mitochondrial function by regulating mitophagy through caveolin-3

**DOI:** 10.1101/2020.10.02.323410

**Authors:** JY Beak, HS Kang, W Huang, A Aghajanian, K Gerrish, AM Jetten, BC Jensen

## Abstract

Preserving optimal mitochondrial function is critical in the heart, which is the most ATP-avid organ in the body. Recently, we showed that global deficiency of the nuclear receptor RORα in the “staggerer” (RORα^sg/sg^) mouse exacerbates angiotensin II-induced cardiac hypertrophy and compromises cardiomyocyte mitochondrial function. The mechanisms underlying these observations have not been defined. Here we present evidence that RORα regulates cardiomyocyte mitophagy using pharmacological and genetic gain- and loss-of-function tools, including a novel cardiomyocyte-specific RORα knockout mouse. Cardiomyocyte RORα is upregulated by hypoxia and the loss of RORα blunts hypoxia-induced mitophagy and broadly compromises mitochondrial function. We show that RORα is a direct transcriptional regulator of the mitophagy mediator caveolin-3 in cardiomyocytes and that increased expression of RORα increases caveolin-3 abundance and enhances mitophagy. Knockdown of RORα impairs cardiomyocyte mitophagy, but this defect can be rescued by caveolin-3 overexpression. Collectively, these findings reveal a novel role for RORα in regulating mitophagy through caveolin-3 and expand our currently limited understanding of the mechanisms underlying RORα-mediated cardioprotection.

## Introduction

The heart consumes more ATP than any other organ by virtue of the requirement for constant cardiomyocyte contraction and relaxation. As such, maintenance of a large and optimally functional pool of mitochondria is essential to normal cardiac physiology. Compromised mitochondrial function leads to energy deprivation and enhanced oxidative stress, both of which are central to the pathobiology of heart failure (Zhou & Tian, 2018). The cardiac mitochondrial pool is preserved through careful orchestration of mitochondrial biogenesis and the clearance of dysfunctional mitochondria, chiefly through selective mitochondrial autophagy, or mitophagy (Kubli & Gustafsson, 2012). Nuclear receptors increasingly are recognized as central transcriptional regulators of these critical processes in the heart (Vega & Kelly, 2017).

The retinoic acid-related orphan nuclear receptor (ROR) subfamily consists of three members, RORα, RORβ, and RORγ (*NR1F1-3*), with highly tissue-specific and context-dependent expression. RORα plays well-described roles in regulating circadian rhythm, lipid metabolism, and T cell development (Cook *et al*, 2015). However, there was no recognized function for RORα in the heart prior to two recent reports from Pu and colleagues, who showed that RORα expression is protective in myocardial ischemia/reperfusion and diabetic cardiomyopathy (He *et al*, 2016; Zhao *et al*, 2017). We extended those findings in the angiotensin II (Ang II)-induced cardiac injury model, demonstrating that “staggerer” mice that globally lack functional RORα (RORα^sg/sg^) develop exaggerated myocardial hypertrophy and contractile dysfunction after Ang II exposure (Beak *et al*, 2019). Using both *in vivo* and *in vitro* approaches we made the novel observation that cardiomyocyte RORα deficiency was associated with decreased mitochondrial abundance, ATP depletion, and enhanced oxidative stress. The mechanisms underlying these findings and putative functions for RORα in uninjured cardiomyocytes remain unclear.

Here we use *in vivo* and *in vitro* approaches to identify a novel role for RORα in regulating cardiomyocyte mitophagy. The absence of RORα is associated with abnormal mitochondrial structure and function, due at least in part to defects in mitophagy mediated by impaired transcriptional regulation of caveolin-3. These new findings may account for our previous observation of energy deprivation in the hearts of RORα^sg/sg^ mice and emphasize the emerging importance of RORα as a cardioprotective nuclear receptor.

## Results

### Mitochondria in RORα^sg/sg^ mouse hearts are less abundant and structurally abnormal with fewer mitophagic events

Our recent study demonstrated that RORα protects cardiomyocytes against Ang II-induced hypertrophy by maintaining mitochondrial number and function (Beak *et al.*, 2019). To explore the potential roles for RORα in preserving the cardiomyocyte mitochondrial pool, we examined wild type (WT) and RORα^sg/sg^ hearts using transmission electron microscopy (TEM). We found that the mitochondria in RORα^sg/sg^ hearts were 44% less abundant but 43% larger than in hearts from WT littermates (**Figures 1A and B**), broadly suggestive of defects in mitochondrial quality control. We also analyzed the TEM images for mitochondria engulfed in double membrane lysosomes, a finding that is indicative of active autophagy, and found 68% fewer of these structures in the RORα^sg/sg^ than the WT hearts. (**Figures 1C and 1D**). Taken together, these findings suggest that defects in mitophagy may contribute to mitochondrial depletion and dysfunction in the absence of RORα.

**Figure 1.**
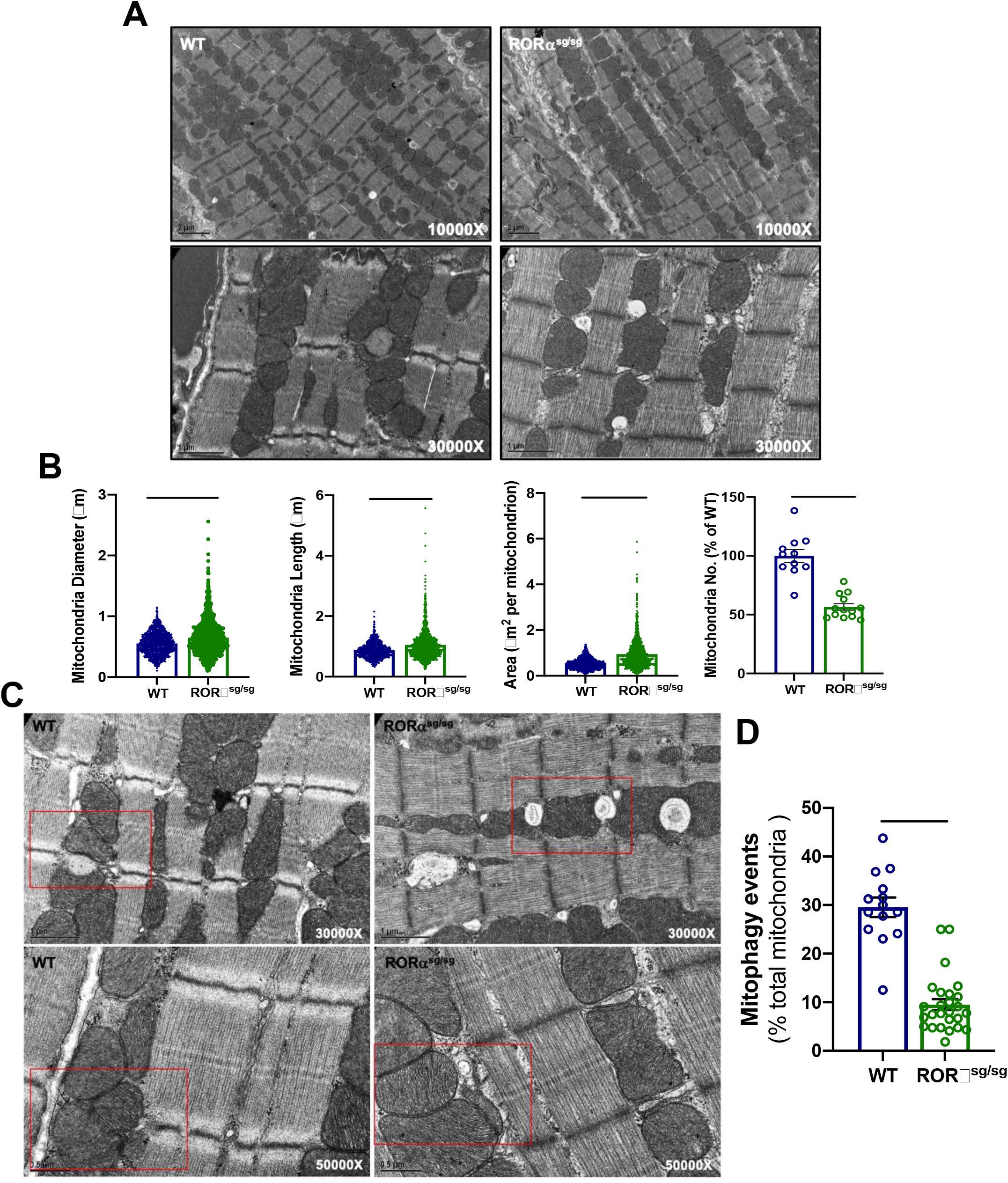
**(A)** Transmission electron microscopy (TEM) of WT and RORα^sg/sg^ hearts; **(B)** Quantitative analysis of TEM mitochondrial abundance and size using Image J with number of mitochondria analyzed (n=2 hearts per group). **(C)** Representative TEM image of mitochondria undergoing lysosomal degradation in WT and RORα^sg/sg^ hearts; **(D)** Quantitation of mitochondria undergoing lysosomal degradation using Image J. **** p <0.0001

### Gene expression profiling suggests a defect in mitochondrial quality control in RORα^sg/sg^ mouse hearts

To identify potential regulation of mitochondrial abundance and function by RORα, we performed microarray analysis using Agilent whole genome mouse oligo arrays. We identified 9653 genes that were differentially expressed in RORα^sg/sg^ (n=4) and WT littermate (n=5) hearts. Of these genes, 3118 were >2.5-fold more abundant and 3340 were >2.5-fold less abundant in RORα^sg/sg^ compared with WT hearts.

To analyze these microarray data for regulation of biologically meaningful pathways, we used the online Database for Annotation, Visualization and Discovery (DAVID) v6.8 (Huang da *et al*, 2009a, b). Filtering for transcripts that were at least 5-fold less abundant in RORα^sg/sg^ hearts revealed a surprising number of pathways related to protein stabilization and degradation (**Figure 2A**). Transcripts in the Gene Ontology term “Protein Stabilization/Lysosomal protein stabilization” (“Any process involved in maintaining the structure and integrity of a protein and preventing it from degradation or aggregation”) were 2.9-fold less abundant in RORα^sg/sg^ hearts compared to WT hearts (p = 1.2 × 10-7, **Figure 2B**). Numerous other significantly affected pathways (**Supplemental Table 1**) were either predicted by the known function of RORα in transcriptional regulation (“DNA templated transcription”, “mRNA processing”) or consistent with our previously published work (Beak *et al.*, 2019) (“Sarcomere organization”, “muscle contraction”, “positive regulation of IκB kinase/NFκB signaling”). Focused analysis of the microarray data demonstrated that the expression of numerous genes associated with mitochondrial function, ATP synthase subunits and autophagy were significantly decreased in the RORα^sg/sg^ mice compared with in WT (**Figure 2C**). Collectively our microarray analysis supported our published data indicating a role for RORα in mitochondrial maintenance and suggested that defects in protein quality control might contribute to mitochondrial dysfunction in RORα^sg/sg^ mouse hearts.

**Figure 2.**
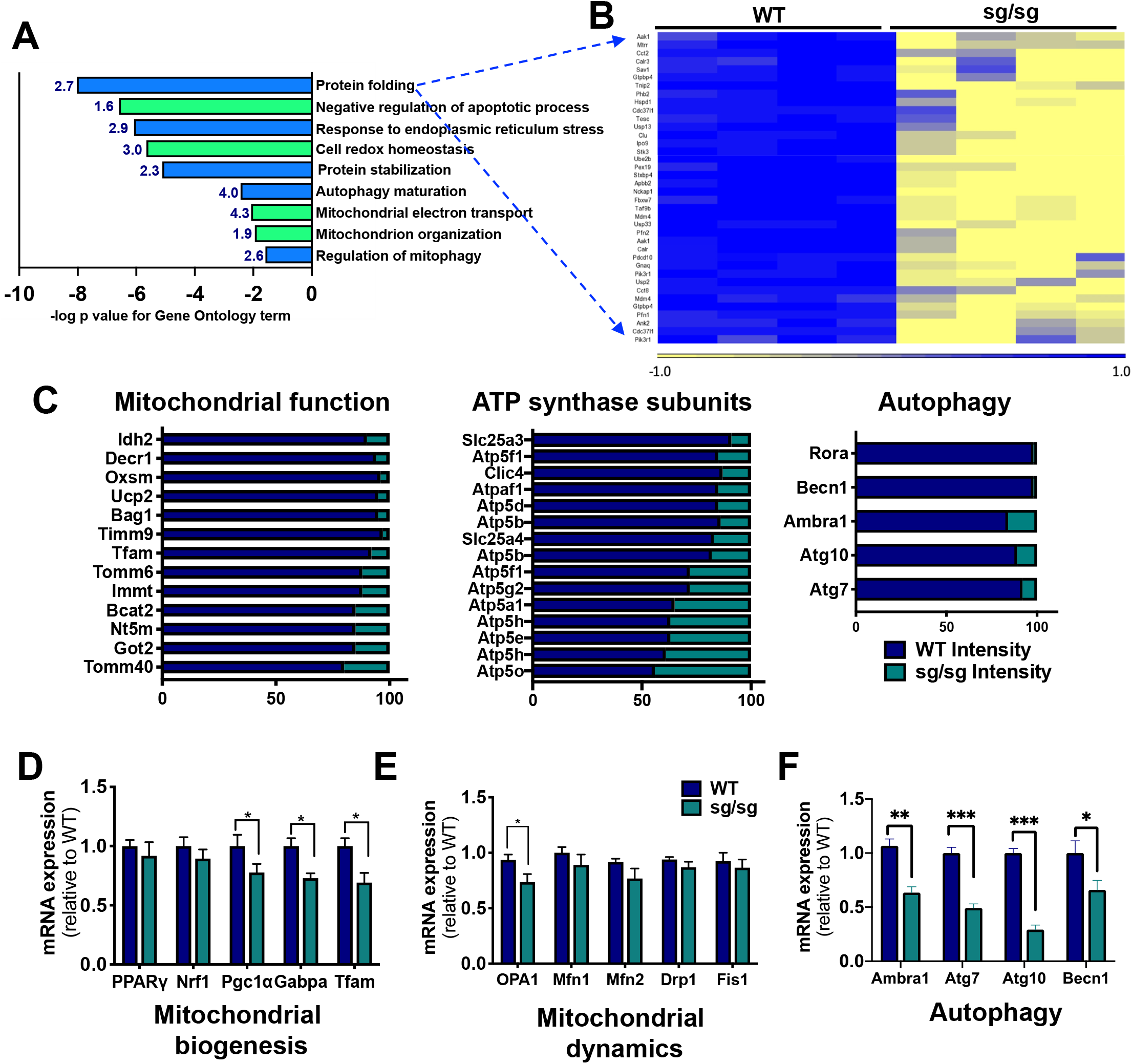
**(A)** DAVID Pathway analysis of RNA microarray data comparing WT (n=5) and RORα^sg/sg^ mouse (n=4) heart with Gene Ontology (GO) pathways and fold-enrichment; **(B)** Heatmap of genes in the GO term Protein Folding; **(C)** Microarray abundance of selected genes related to mitochondrial function, ATP synthase subunits, and autophagy; **(D-F)** Quantitative reverse transcriptase PCR for selected genes (n=3-6 mice per group). Ambra1 = autophagy and beclin 1 regulator 1; Atg7 and 10 = autophagy-related protein 7 and 10; Atp5f = ATP synthase F(0) complex subunit B1; Atp5g1, 2 and 3 = ATP synthase, H+ transporting, mitochondrial Fo complex subunit C1, C2, and C3; Atp5k = ATP synthase membrane subunit e, Atp5o; ATP synthase, H+ transporting; Cox6b = cytochrome c oxidase subunit 6b; Drp1 = Dynamin related protein 1; Fis1 = mitochondrial fission 1 protein; LC3 = microtubule-associated protein 1A/1B-light chain 3; Mfn1 and Mfn2 = Mitofusin 1 and 2; Ndufs2 and 4, Ndufv1 = NADH Ubiquinone oxidoreductase core subunit S2 and S4; OPA1 = mitochondrial dynamin like GTPase; Pink1 = PTEN-induced kinase 1. * p <0.05, ** p <0.01, *** p <0.001

To further explore at a transcriptional level whether alterations in mitochondrial biogenesis, dynamics, or autophagy might determine the compromised mitochondrial phenotype in the RORα^sg/sg^ mouse hearts, we performed quantitative reverse transcriptase polymerase chain reaction (qRT-PCR) for selected genes critical to each process. Abundance of *PGC1α*, *Gabpa*, and *TFAM* (mitochondrial biogenesis, **Figure 2D**) was mildly reduced, as was expression of *Opa1* (mitochondrial dynamics, **Figure 2E**). However, the most profound transcriptional differences were found in the autophagy-related genes autophagy and beclin 1 regulator 1 (*Ambra1*), autophagy related 7 (*Atg7*), *Atg10*, and beclin1 (*Becn1*) (**Figure 2F**). Using qRT-PCR, we also confirmed decreased abundance of multiple key electron transport genes, further corroborating the presence of a RORα-related mitochondrial defect (**Supplemental Figure S1**). These findings coupled with the morphological abnormalities seen on TEM (**Figure 1**) and transcriptome pathway analysis (**Figure 2A**) encouraged us to determine whether and how RORα regulates cardiomyocyte mitophagy.

### RORα regulates expression of mitophagy mediators in normoxic and hypoxic conditions *in vivo* and *in vitro*

Mitophagy is carried out primarily through two well-defined pathways: PTEN-induced putative kinase 1 (PINK1) /Parkin and BCL2 interacting protein 3 (Bnip3)/NIX. Activation of either of these pathways can target mitochondria for autophagy by linking to microtubule associated protein light chain beta 3-II (LC3B-II) on autophagosome membranes (Moyzis *et al*, 2015). To better understand how RORα might affect mitophagy, we immunoblotted the mitochondrial fraction of WT and RORα^sg/sg^ mouse hearts for these canonical mitophagy mediators and found that the absence of functional RORα was associated with markedly decreased abundance of mitochondrial Bnip3, PINK1, and LC3B-II (**Figure 3A**).

**Figure 3.**
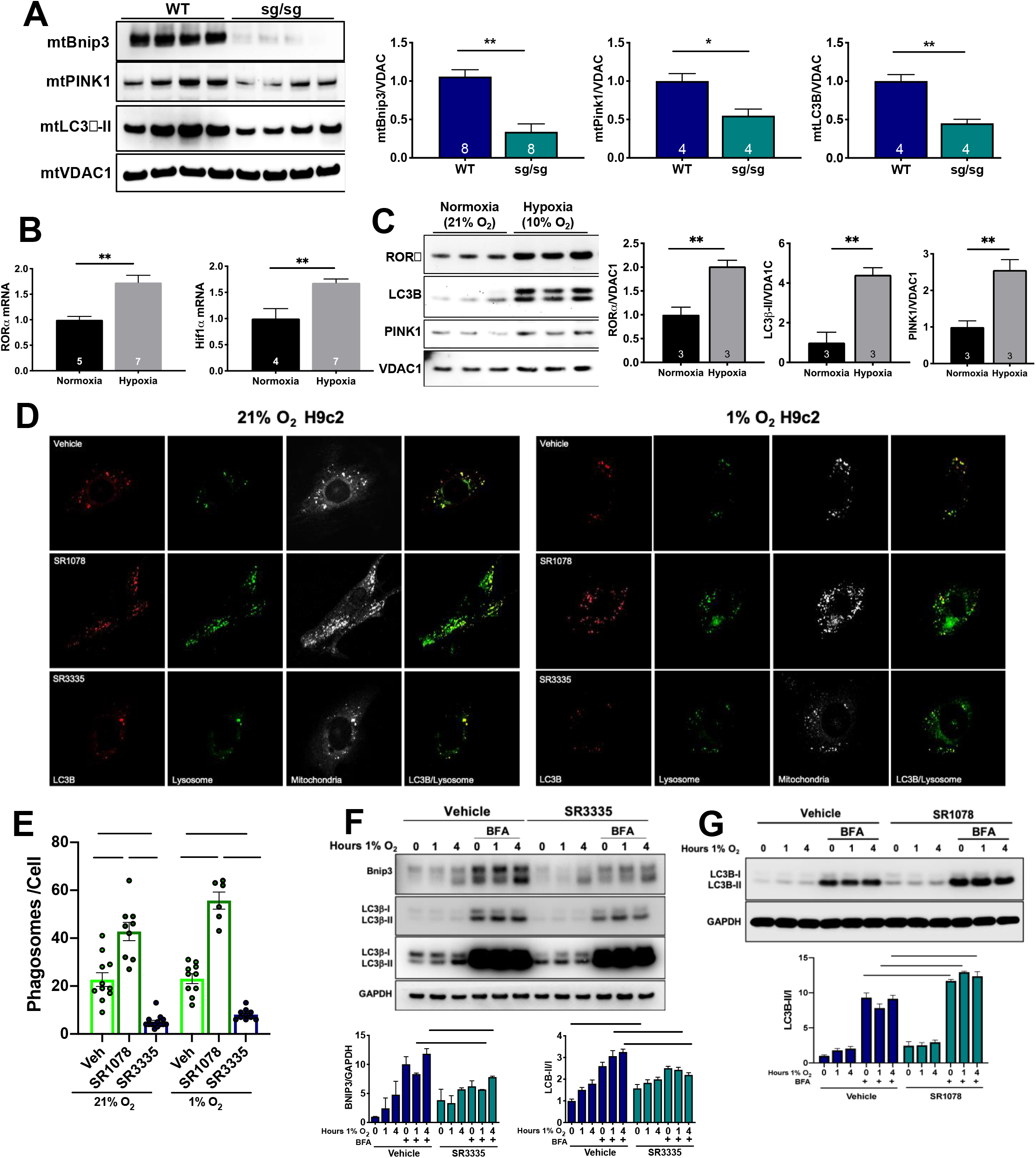
**(A)** Immunoblotting of mitochondrial fraction from WT and sg/sg heart lysates. **(B)** Quantitative reverse transcriptase PCR on hearts of mice exposed to normoxia or hypoxia (10% O_2_) for 24 hr; **(C)** Immunoblotting of heart lysates of mice exposed to normoxia or hypoxia. **(D)** Immunofluorescent staining of normoxic or hypoxic (1% O_2_ for 24 hr) H9c2 cardiomyoblasts exposed to vehicle, RORα agonist SR1078 (10μM), or RORα inverse agonist SR3335 (20μM). **(E)** Quantitative analysis of phagosome indexed by co-staining for LC3B, the lysosomal stain LysoTracker, and mitochondria using Image J. **(F)** Immunoblotting of NRVM lysates exposed to hypoxia in the presence or absence of bafilomycin A with SR3335 (20μM) or **(G)** SR1078 (10μM) with summary densitometry (Image J); BFA = bafilomycin A; Bnip3 = BCL2 interacting protein 3; Hif1α = Hypoxia inducible factor1 alpha; LC3 = microtubule-associated protein 1A/1B-light chain 3; Pink1 = PTEN-induced kinase 1; VDAC1 = Voltage Dependent Anion Channel 1. * p <0.05, ** p <0.01, *** p <0.001, **** p <0.0001

Oxygen tension is a potent regulator of mitochondrial function in the heart and protecting mitochondrial function is critical when the myocardium becomes ischemic (Cadenas *et al*, 2010). Under hypoxic conditions, such as myocardial infarction, mitophagy is upregulated to promote clearance of dysfunctional mitochondria. Failure to induce mitophagy can lead to exacerbated cardiomyocyte injury and impaired survival (Kubli *et al*, 2013). To assess the *in vivo* function of RORα in low oxygen tension, we subjected WT mice to hypoxia (10% O2 for 24 hours). Hypoxia led to nearly 2-fold upregulation of myocardial RORα mRNA expression, similar in magnitude to the classical hypoxia marker, Hif1α (**Figure 3B**). Immunoblots of heart lysates revealed that hypoxic upregulation of RORα was coupled with significant increases in LC3B and PINK1 expression, consistent with induction of mitophagy (**Figure 3C**).

Next we sought to determine whether pharmacological manipulation of RORα activity *in vitro* could recapitulate our genetic loss-of-function findings from our *in vivo* models. We exposed H9c2 rat ventricular myoblasts to hypoxia (1% O_2_) in the presence and absence of the RORα inverse agonist SR3335 or agonist SR1078 then quantified the number of phagolysosomes co-stained for LC3B (red) and the lysosomal stain LysoTracker (green) (**Figure 3D**). The RORα agonist SR1078 increased and the RORα inverse agonist SR3335 decreased phagolysosome events in both normoxia and hypoxia (**Figure 3E**). We then exposed normoxic and hypoxic neonatal rat ventricular myocytes (NRVMs) treated with SR3335 or SR1078 to bafilomycin A1 (BFA), which interferes with autolysosome acidification and autophagosome formation through inhibition of V-ATPase (Mauvezin & Neufeld, 2015), allowing measurement of autophagic flux as indicated by the ratio of LC3B-II/I (Sharifi *et al*, 2015). BFA increased the expression of the mitophagy adaptor Bnip3, as well as the LC3B-II/I ratio as expected in vehicle-treated NRVMs. SR3335 abrogated upregulation of Bnip3 and blunted the LC3B-II/I increase (**Figure 3F**) whereas SR1078 increased LC3B-II (**Figure 3G**), suggesting that RORα regulates hypoxic autophagic flux

Collectively, these findings suggest that upregulation of RORα in response to hypoxia facilitates cytoprotective autophagy, corroborating our morphological (**Figure 1**) and transcriptional (**Figure 2)** findings.

### Cardiomyocyte mitophagy is compromised *in vivo* in a novel cardiomyocyte-specific RORα knockout mouse

The RORα^sg/sg^ “staggerer” mouse is a well-validated genetic loss-of-function model, but it exhibits numerous extracardiac phenotypes that could potentially influence our study of mitophagy in the heart. Though our *in vitro* studies support cardiomyocyte autonomous functions for RORα, we created a cardiomyocyte-specific RORα knock-out (CMKO) mouse, breeding floxed RORα mice with αMHC-Cre mice, to extend those findings *in vivo* and to eliminate potential contributions from extracardiac phenotypes. Cre mRNA was detected only in the heart and RORα mRNA abundance was 80% lower in the CMKO than CMWT (*Cre*^*−*^*flox*^*+/+*^) hearts, expression in liver and kidney was unaffected, validating our breeding strategy. We found no compensatory changes in RORγ transcriptional expression (**Figure 4A**).

**Figure 4.**
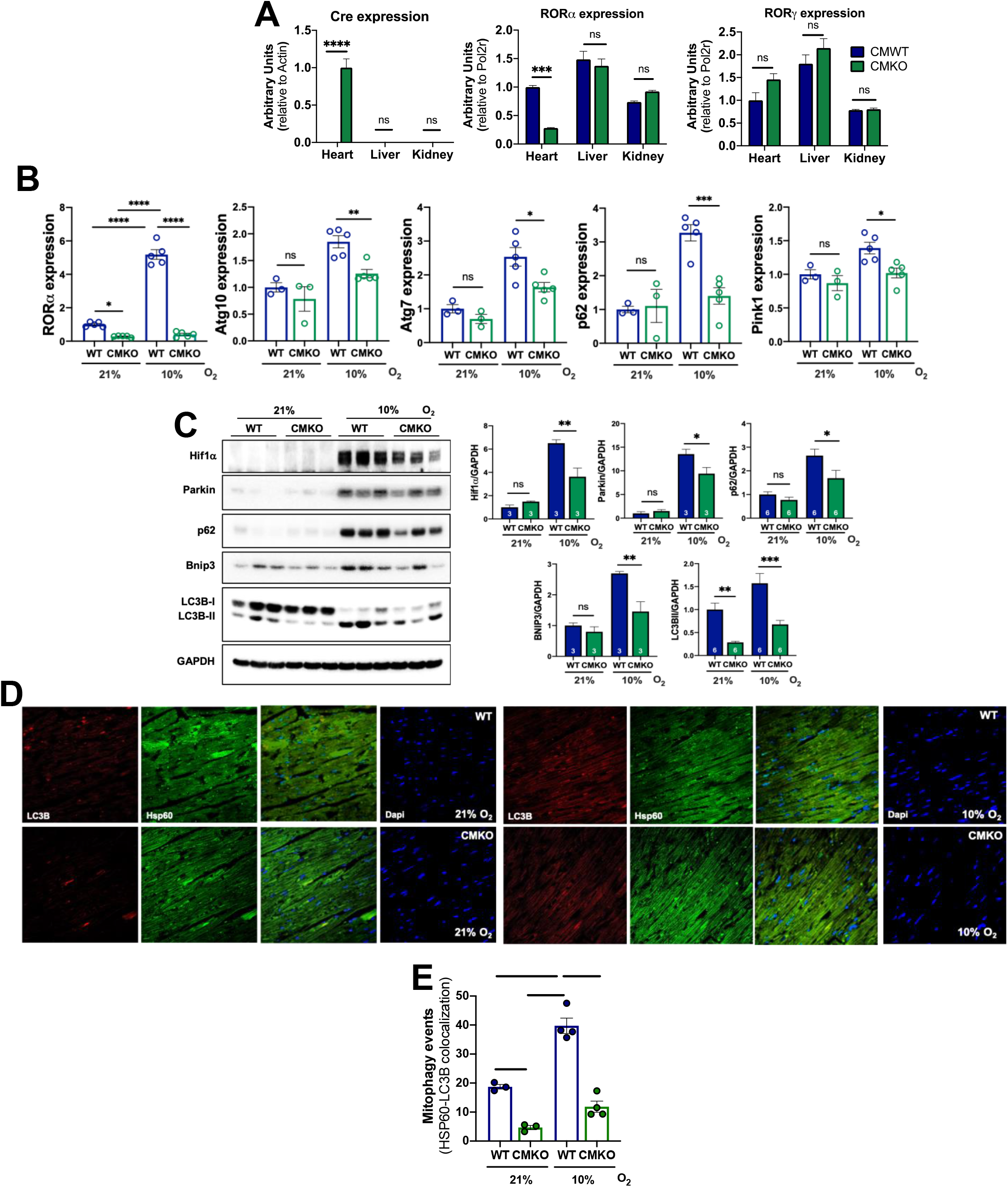
**(A)** Quantitative reverse transcriptase PCR (qRT-PCR) for Cre, RORα and γ genes on heart, liver, and kidney from CMWT and cardiomyocyte specific RORα knock out (CMKO) mice (n=3 per group). **(B)** qRT-PCR for autophagy-inducing genes in heart tissue from CMWT and CMKO mice exposed to normoxia (21% O_2_) or to hypoxia (10% O_2_) for 8 hrs. **(C)** Immunoblotting from WT and CMKO heart lysates with summary of densitometry (Image J). **(D, E)** Representative immunofluorescence images of mouse heart sections and Image J summary quantification; Atg10 = autophagy related 10; Atg7 = autophagy related 7; Bnip3 = BCL2 interacting protein 3; Cre = Cre recombinase; DAPI = 4′,6-diamidino-2-phenylindole; GAPDH = Glyceraldehyde 3-phosphate dehydrogenase; Hif1α = Hypoxia inducible factor1 alpha; Hsp60 = Heat shock protein 60; LC3B = microtubule-associated protein 1A/1B-light chain 3; PINK1 = PTEN-induced kinase 1; p62 = SQSTM1; VDAC1 = Voltage Dependent Anion Channel 1. * p <0.05, ** p <0.01, *** p <0.001, **** p <0.0001

We then examined the effect of cardiomyocyte-specific RORα deletion on the *in vivo* expression of several proteins that are centrally involved in the induction and maintenance of mitophagy. RNA was isolated from hearts of CMWT and CMKO mice that were housed in normoxic (21% O_2_) or hypoxic (10% O_2_) conditions for 8 hours. Abundance of the transcripts encoding Atg10 and Atg7, autophagy receptor p62, and PINK1 was higher in hypoxic than normoxic CMWT hearts. However, the hypoxia-induced increase in these critical autophagy mediators was not observed in CMKO mouse hearts (**Figure 4B**). Immunoblotting demonstrated the expected increases in Parkin, p62, Bnip3, and LC3B-II in hypoxic CMWT hearts but these changes were blunted in CMKO hearts (**Figure 4C**). We next sought to examine the functional consequences of these differences, using immunofluorescent staining (**Figure 4D**) for LC3B (red) and the mitochondria marker heat shock protein-60 (Hsp60, green). In normoxic conditions, CMKO hearts exhibited 75% fewer mitophagy events than CMWT hearts (defined by LC3B and Hsp60 co-staining). Hypoxia induced an increase in mitophagy in both CMWT and CMKO hearts, but mitophagosomes were 70% less abundant in CMKO than CMWT (**Figure 4E**).

Collectively, these results substantiate our findings that RORα regulates mitophagy in the heart. They further indicate that the deficits in mitophagy exhibited by the global RORα^sg/sg^ mouse most likely arise from cardiomyocyte autonomous effects of RORα.

### Loss of RORα compromises hypoxic mitophagy and mitochondrial function *in vitro*

The previous findings encouraged us to further analyze the effect of RORα on mitophagy and mitochondrial function in hypoxic cardiomyocytes. To measure mitophagy in neonatal rat ventricular myocytes (NRVMs), we used mitochondrial-targeted Keima (mt-Keima), a fluorescent protein pH indicator that fluoresces red (543 nm) when introduced to the acidic environment of the lysosome but green (458 nm) at normal or basic pH (Shirakabe *et al*, 2016).

We infected NRVMs with lentivirus containing either short hairpin RNA (shRNA) against RORα or scrambled shRNA (shCtrl) and found that the mt-Keima fluorescence ratio (543/458 nm) underwent the expected increase under hypoxic conditions but was decreased in both normoxia and hypoxia by RORα knockdown (**Figure 5A**).

**Figure 5.**
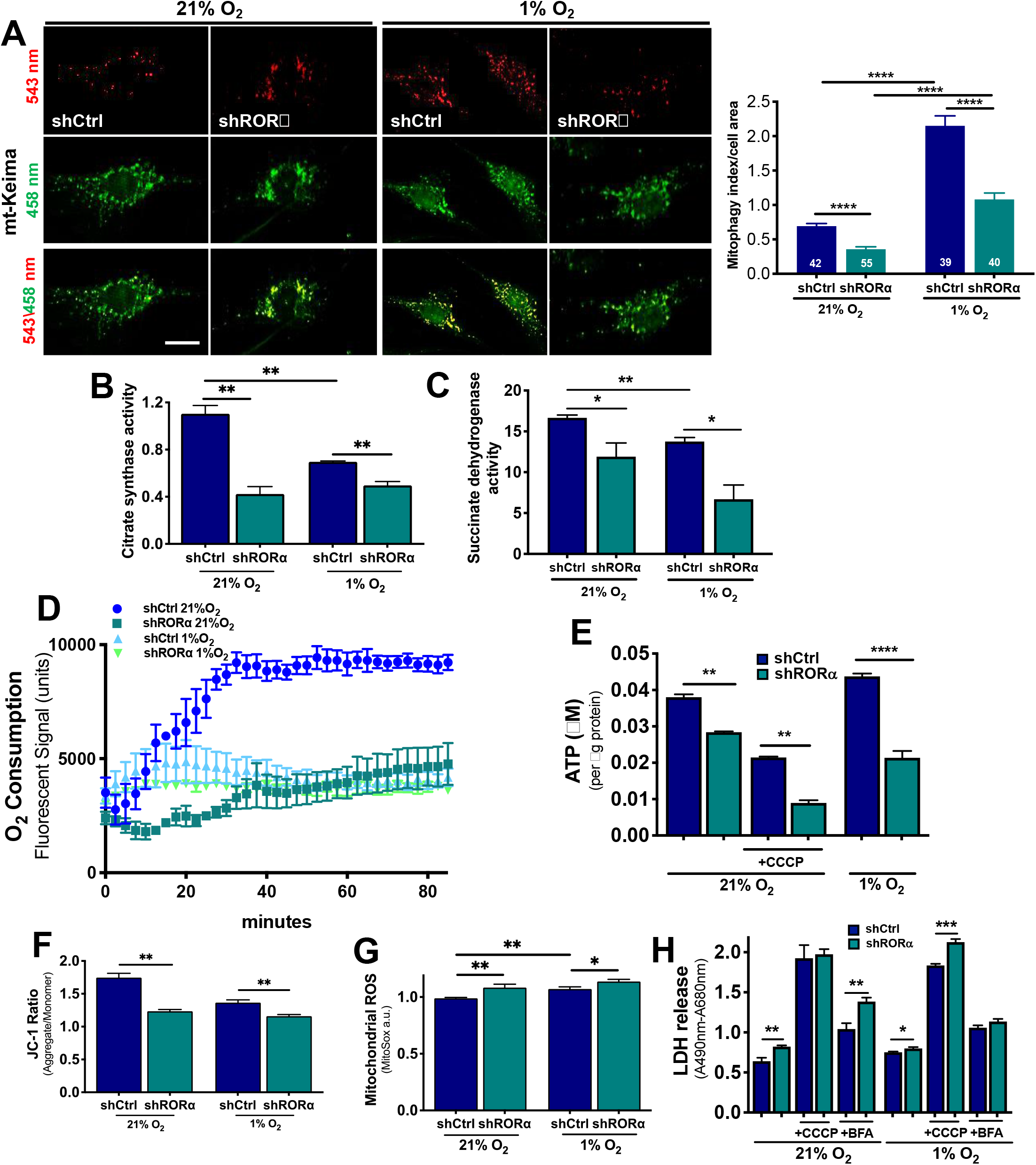
**(A)** Representative immunofluorescent images of NRVMs infected with the mitophagy indicator, mitochondria specific Keima (mtKeima) for 1hr with summary fluorescent quantification using Image J; **(B)** Citrate synthase and **(C)** succinate dehydrogenase activity in NRVMs infected with lentivirus containing scrambled shRNA (shCtrl) or shRNA against RORα (shRORα) and exposed to normoxia (21%) and hypoxia (1%) for 24hr; **(D)** Oxygen consumption rate in NRVMs infected with lentivirus containing shCtrl or shRORα then exposed to normoxia or hypoxia; **(E)** ATP concentration in NRVMs infected shCtrl or shRORα in the presence or absence of carbonyl cyanide m-chlorophenyl hydrazine (CCCP), a potent uncoupler of mitochondrial oxidative phosphorylation; **(F)** NRVM mitochondrial membrane potential assayed using JC-1 (5, 5′, 6, 6′-tetrachloro-1, 1′, 3, 3′-tetraethylbenzimidazolylcarbocyanine iodide). Summary fluorescence used Image J; (**G**) Mitochondrial reactive oxygen species (ROS) assayed using MitoSox with summary fluorescence using Image J; **(H)** Cell death assayed by LDH release in NRVMs exposed to normoxia or hypoxia in the presence or absence of bafilomycin A (BFA) or CCCP. * p <0.05, ** p <0.01, *** p <0.001, **** p <0.0001

To expand upon our earlier *in vivo* discovery that RORα^sg/sg^ mouse hearts have decreased ATP abundance (Beak *et al.*, 2019), we used lentiviral shCtrl or shRORα in NRVMs for an *in vitro* loss-of-function approach. Knockdown of RORα in NRVMs decreased the functional mitochondrial mass, as measured by citrate synthase (**Figure 5B**) and succinate dehydrogenase (**Figure 5C**) activity. Oxygen consumption rate was markedly decreased by shRORα (**Figure 5D**). These metabolic insults collectively contributed to decreased ATP abundance in the absence of functional RORα (**Figure 5E**). Knockdown of RORα compromised mitochondrial membrane potential as measured by JC-1 (**Figure 5F**) and increased mitochondrial reactive oxygen species (ROS) generation as assayed by MitoSox (**Figure 5G**). Cell death, as measured by LDH release, was only modestly increased by shRORα (**Figure 5H**). Taken together, these findings further indicate that RORα contributes to optimal mitochondrial function in both normoxia and hypoxia, likely through maintenance of mitophagy.

### RORα directly binds the caveolin-3 promoter to regulate hypoxic mitophagy

Caveolin-3 (Cav-3) enhances autophagy and protects mitochondrial function in hypoxic HL-1 cardiomyoblasts (Kassan *et al*, 2016). Hence, we hypothesized that RORα promotes mitophagy in the heart by regulating the expression of Cav-3.

Analysis of Cav-3 protein demonstrated that the level of Cav-3 protein was 34% lower in RORα^sg/sg^ than WT hearts (**Figure 6A**). In the cardiomyocyte-specific RORα CMKO, Cav-3 mRNA (**Figure 6B**) and protein (**Figures 6C and 6D**) levels were similar to WT hearts in normoxic conditions. After WT and CMKO mice spent 8 hours in a hypoxia chamber (10% O_2_), Cav-3 mRNA and protein were increased 2-3-fold in WT hearts, but there was no change in Cav-3 abundance in CMKO hearts. These findings suggested that RORα may regulate Cav-3 expression in cardiomyocytes. Through promoter analysis, we found two consensus RORα binding sites (ROREs) in the proximal promoter of Cav-3. To determine whether the *Cav3* gene is a direct transcriptional target gene of RORα in adult mouse ventricular myocytes, we performed chromatin immunoprecipitation (ChIP)-PCR using immunopulldown with an anti-RORα antibody or an anti-IgG antibody as a negative control. We found significant enrichment of the regions containing the two ROREs (−0.1 kb, and −0.5 kb regions), but not in a region of the promoter that does not contain an RORE (−4.2kb), indicating that RORα regulates *Cav3* directly (**Figure 6E**).

**Figure 6.**
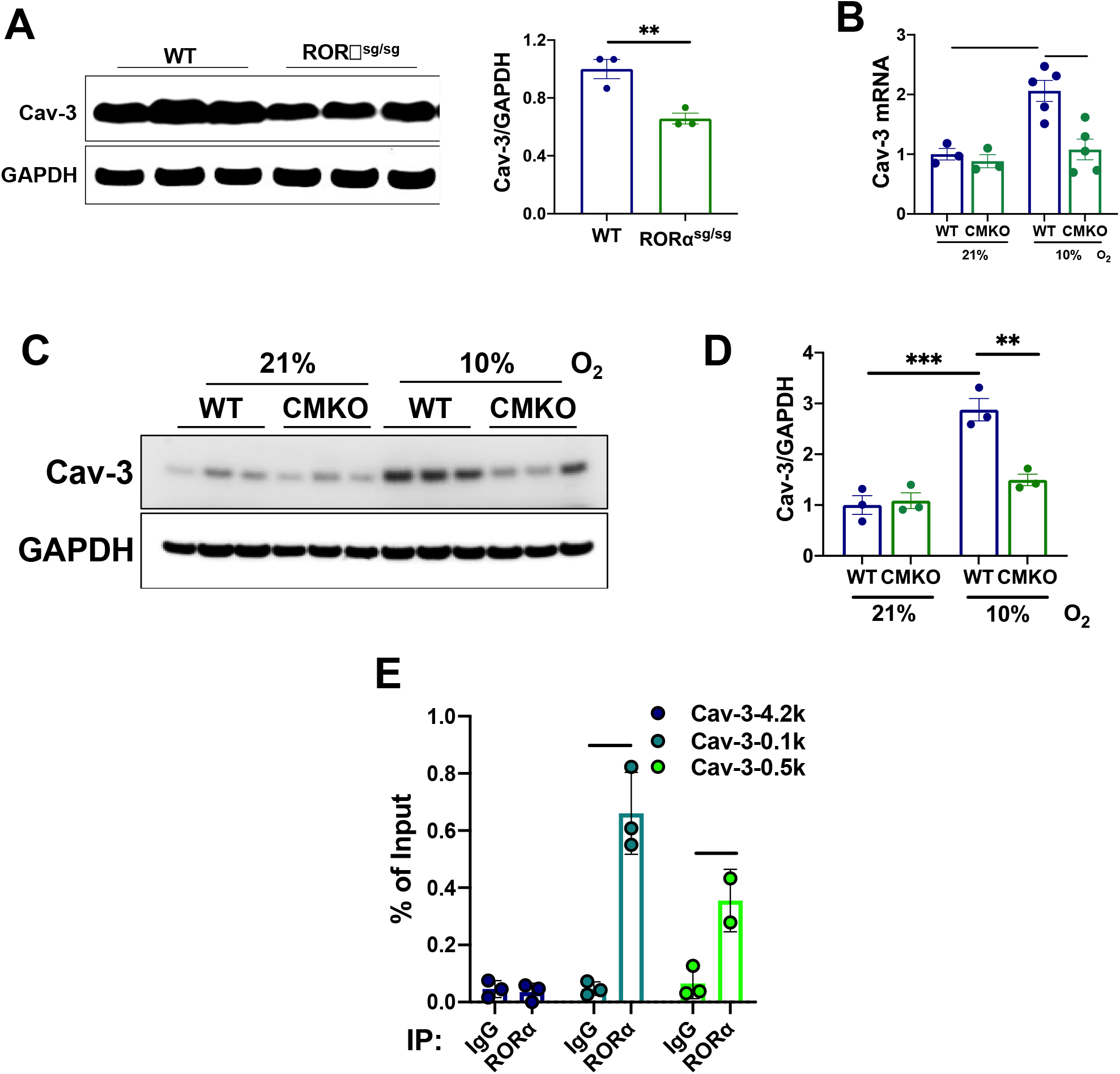
**(A)** Immunoblotting of heart lysates from WT and sg/sg mouse hearts with summary densitometry (Image J); **(B)** Quantitative reverse transcriptase PCR for Cav-3 in hearts of WT and CMKO mice exposed to nomoxia (21% O_2_ for 8 hrs) or hypoxia (10% O_2_ for 8 hrs). **(C)** Immunoblotting from WT and CMKO heart lysates with **(D)** summary densitometry (Image J); **(E)** Chromatin immunoprecipitation (ChIP)-PCR of Cav-3 in adult mouse ventricular myocytes incubated with anti-RORα or anti-IgG (control) antibodies. Cav-3 = Caveolin-3; GAPDH = Glyceraldehyde 3-phosphate dehydrogenase; IP = Immuno-pull down. * p <0.05, ** p <0.01, *** p <0.001, **** p <0.0001

Next, we examined the effect of RORα expression on Cav-3 expression in H9c2 cardiomyoblasts infected with a lentivirus containing doxycycline-inducible RORα4 (H9c2-Flag-RORα4-HA), the predominant RORα isoform in the heart (**Supplemental Figure S2A**). Doxycycline induced RORα4 in a dose-dependent manner (**Supplemental Figure S2B**) and RORα4 was localized largely to nuclei, as expected (**Supplemental Figure S2C**). The induction of RORα by doxycycline resulted in a 4.5-fold increase in Cav-3 expression (**Figure 7A**). The expression of Cav-3 was then compared in doxycycline-treated and untreated H9c2-Flag-RORα4-HA cells exposed to either hypoxia (1% O_2_) or normoxia (21% O_2_). We found that hypoxia enhanced Cav-3 expression, and this upregulation was significantly greater in doxycycline-treated cells in which RORα was induced (**Figures 7B and 7C).**To determine whether this increase in Cav-3 expression correlated with enhanced mitophagy we quantified co-localization of immunofluorescent LC3B (red) with mitochondria (green) and identified roughly 4 times more mitophagy events in the cells in which RORα was induced by doxycycline (**Figure 7D**). To determine whether Cav-3 is directly involved in the regulation of cardiomyocyte mitophagy by RORα we knocked down RORα in H9c2 cardiomyoblasts with shRNA then transfected these cells with an EGFP-Cav-3 plasmid. We then visualized mitophagy events by co-staining for LC3B (blue) and mitochondria (red), using EGFP to identify Cav-3 (**Figure 7E**). As expected, shRORα decreased mitophagy events, however forced expression of Cav-3 eliminated this defect such that mitophagy events were equivalent in shCtrl-Cav-3+ and shRORα-Cav3+ H9c2s. These results support the hypothesis that RORα directly regulates the expression of Cav-3 in cardiomyocytes and that RORα-mediated Cav-3 induction promotes mitophagy. Taken together, these findings provide a potential mechanism for the mitochondria-protective effects of RORα in the heart.

**Figure 7.**
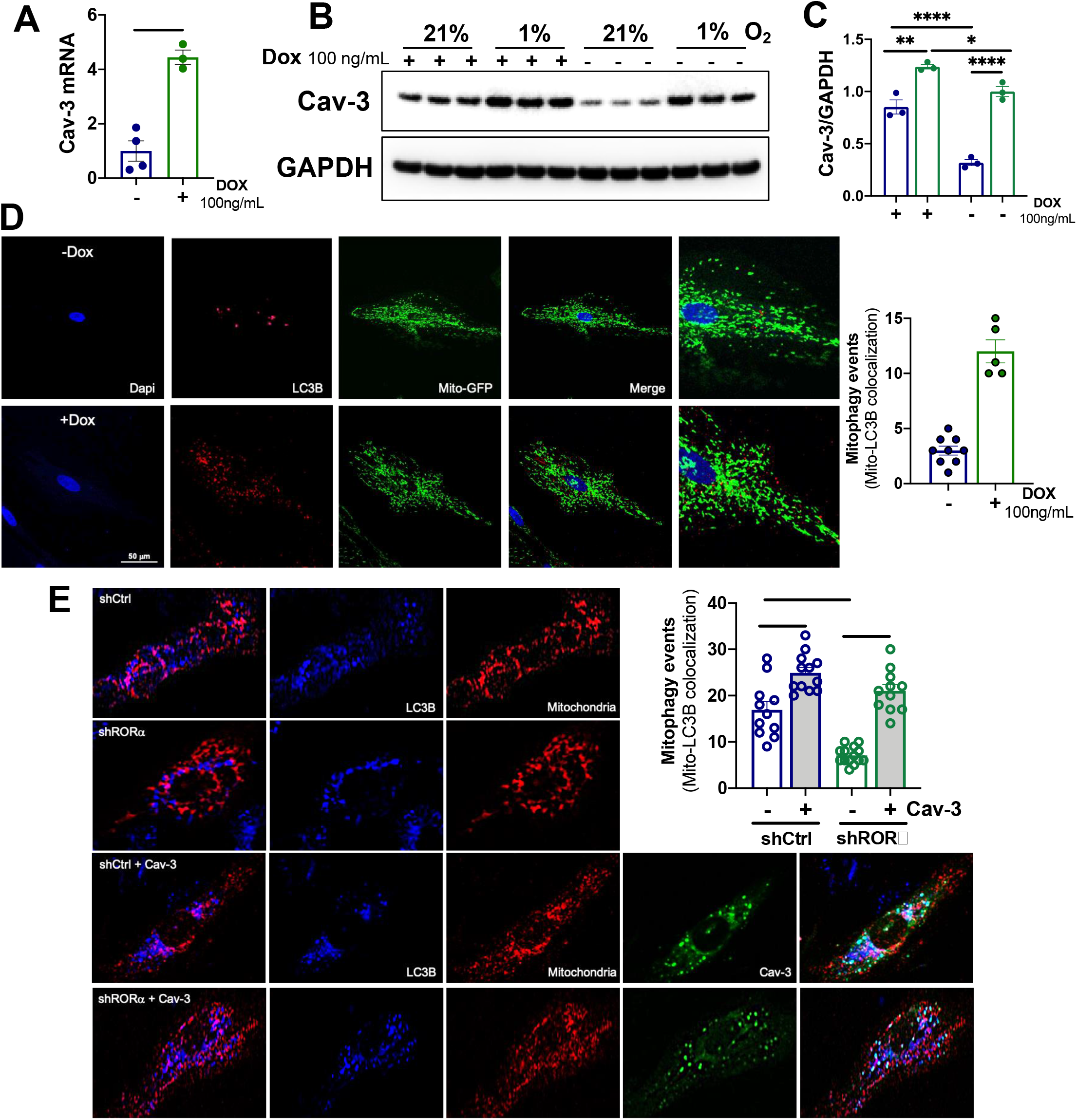
**(A)** Quantitative reverse transcriptase PCR for Cav-3 in H9c2-Flag/RORα4/HA cells. **(B)** Immunoblotting of cell lysates with **(C)** summary densitometry from H9c2-Flag/RORα4/HA cultured in normoxia (21% O_2_) or hypoxia (1% O_2_) for 24 hrs in the absence or presence of doxycycline induction of RORα. **(D)** Immunofluorescence staining of H9c2-Flag/RORα4/HA cells with summary quantitation of LC3B and Mito-GFP costaining (Image J); (E) H9c2-Flag/RORα4/HA cells with m-Raspberry-mito-7 with or without EGFP-Cav-3 plasmid transfection. Cav-3 = Caveolin-3; DAPI = 4′,6-diamidino-2-phenylindole; Dox = Doxycycline; GAPDH = Glyceraldehyde 3-phosphate dehydrogenase; LC3 = microtubule-associated protein 1A/1B-light chain 3. * p <0.05, ** p <0.01, *** p <0.001, **** p <0.0001

## Discussion

In this study, we identify a novel role for the nuclear receptor RORα in the regulation of mitophagy, in part through transcriptional upregulation of Cav-3. Loss of RORα profoundly compromises cardiomyocyte mitochondrial function leading to energy deprivation, oxidative stress and cell death. These insults are particularly deleterious in the setting of hypoxia. The heart is the most ATP-avid organ in the body, necessitating optimal mitochondrial capacity. Nuclear receptors such as the peroxisome proliferator activated receptor (PPAR) and estrogen related receptor (ERR) families are central regulators of cardiomyocyte mitochondrial structure and function (Vega & Kelly, 2017), and the present findings identify RORα as an additional nuclear receptor that contributes to maintaining cardiac mitochondrial health. A role for RORα in regulating mitochondrial fission in the liver was proposed previously (Kim *et al*, 2017), but to our knowledge our present study is the first to demonstrate that RORα adaptively regulates mitophagy in any tissue.

We chose to use hypoxia as an *in vivo* and *in vitro* stressor to test the role of RORα in the cardiac response to stress because hypoxia is known to stimulate mitophagy and because the cellular response to hypoxia is critical to important clinical conditions such as ischemia-reperfusion injury and myocardial infarction. We found that RORα was upregulated by hypoxia in cardiomyocytes (**Figures 3B and C**) and that RORα knockdown broadly compromised cardiomyocyte mitochondrial function under hypoxic conditions (**Figure 5**). We also show that our novel mice with a cardiomyocyte-specific RORα deletion exhibit a failure to induce hypoxic mitophagy *in vivo* (**Figure 4**). Though the role of RORα in the cardiac response to hypoxia has not been studied previously, a complex interaction has been demonstrated in other tissues between RORα and hypoxia induced factor-1α (HIF-1α), the canonical regulator of cellular response to low oxygen tension. RORα regulates the transcription of hypoxia HIF-1α in endothelial cells (Kim *et al*, 2008). Interestingly, hypoxia increases RORα transcription in a hepatoma cell line by binding to an HRE (hypoxia response element) in the RORα promoter (Chauvet *et al*, 2004), indicating the potential for reciprocal regulation of RORα and HIF-1α. In future studies we will explore whether RORα regulates other critical elements of the cardiomyocyte response to hypoxia.

Our previous work demonstrated that RORα mitigated angiotensin II-induced cardiac hypertrophy through the regulation of cardiomyocyte STAT3 signaling (Beak *et al.*, 2019). Here we identify a novel role for RORα in promoting cardiomyocyte mitophagy through enhancing *Cav3* transcription. Like the other two members of the caveolin family of proteins, Cav-3, the predominant caveolin isoform in muscle cells, serves primarily as a scaffold for membrane trafficking and signaling within caveolae (Tang *et al*, 1996). However, Cav-3 also dissociates from caveolae to the cytosol in injured cardiomyocytes (Ratajczak *et al*, 2003) and to the mitochondrial membrane where it enhances respiratory capacity and mitigates oxidative stress (Fridolfsson *et al*, 2012). Cardiac specific overexpression of Cav-3 protects against hypertrophy induced by pressure loading (Horikawa *et al*, 2011) and ischemia-reperfusion injury (Tsutsumi *et al*, 2008). Cav-3 knockout mice exhibit impaired cardiomyocyte mitochondrial function (Fridolfsson *et al.*, 2012) and enhanced susceptibility to injury (Horikawa *et al*, 2008). Recognized mechanisms for these protective effects include preservation of t-tubular calcium flux(Kong *et al*, 2019) and prevention of hypoxia-induced apoptosis.(Zhou *et al*, 2018)

As a nuclear receptor, RORα is thought to exert most of its effects through transcriptional regulation. RORα regulates caveolin-3 expression by directly binding the *Cav3* promoter in skeletal muscle (Lau *et al*, 2004). The importance of caveolins in regulating autophagy has been established in cardiomyoblasts and other cell types. Cav-3 is upregulated by simulated ischemia-reperfusion in HL-1 cardiomyoblasts and knockdown of Cav-3 in HL-1s compromises autophagy and mitochondrial function, contributing to enhanced cell death.(Kassan *et al.*, 2016) Cav-1 interacts with beclin-1 to induce autophagolysosome formation and protect against cerebral oxidative stress and ischemic injury (Nah *et al*, 2017). Here we show that RORα binds two sites in the *Cav3* promoter to regulate Cav-3 expression (**Figure 6E**). Cav-3 abundance is lower in the hearts of RORα^sg/sg^ mice (**Figures 6A and 6B**) and cardiomyocyte-specific RORα deletion is associated with a failure of the expected hypoxic induction of Cav-3 (**Figures 6C and 6D**). We also demonstrate that RORα overexpression in cardiomyocytes induces Cav-3 expression (**Figures 7A-C**) and promotes mitophagy (**Figure 7D**) in normoxic and hypoxic conditions. Forced expression of Cav-3 rescues mitophagy defects after RORα knockdown in cardiomyocytes (**Figure 7E**). Taken together these findings suggest that RORα and Cav-3 might represent part of a coordinated and protective response to hypoxia in the heart that includes enhanced mitophagy.

We began these investigations to expand upon our earlier finding that the global absence of functional RORα was associated with decreased mitochondrial number and ATP depletion in the heart (Beak *et al.*, 2019). Here we show that knocking down RORα in cardiomyocytes reduces citrate synthase activity and respiratory capacity while compromising mitochondrial membrane potential and increasing oxidative stress, the first detailed characterization of how RORα affects mitochondrial function. Though our central conclusion is that RORα regulates mitophagy to preserve the cardiomyocyte mitochondrial pool, we also present evidence that RORα may be involved in other aspects of mitochondrial homeostasis. We show that the expression of several key regulators of mitochondrial biogenesis, PGC1α, Gabpa, and TFAM, was lower in RORα^sg/sg^ than WT hearts (**Figure 2D**). PGC1α is more abundant in RORα^sg/sg^ liver cells (Lau *et al*, 2008), but less abundant in skeletal muscle cells lacking functional RORα (Lau *et al.*, 2004), indicating that this regulation is tissue-dependent. We also show that Opa1, a canonical regulator of mitochondrial fission, is reduced in RORα^sg/sg^ hearts (**Figure 2E**), consistent with a previous report that RORα regulates mitochondrial dynamics in hepatocytes (Kim *et al.*, 2017). As such, it is possible that induction of mitophagy alone does not account for the protective effects of RORα on mitochondrial function. Indeed, in future studies we will explore potential roles for RORα in the regulation of mitochondrial biogenesis and/or dynamics in cardiomyocytes.

In summary, our experiments indicate that RORα protects cardiomyocyte mitochondrial function through regulation of mitophagy. These findings provide mechanistic underpinning for recent studies demonstrating an emerging cardioprotective role for RORα in the setting of ischemia-reperfusion, diabetic cardiomyopathy and angiotensin II-induced heart failure. The RORα agonist SR1078 recently was shown to be well-tolerated and efficacious in an animal model of autism (Wang *et al*, 2016) and our present study indicates that this agonist is active in normoxic and hypoxic cardiomyocytes (**Figures 3E and G**). Taken together, these studies suggest that activation of RORα *in vivo* could represent a novel therapeutic approach to heart disease.

**Sources of funding: BCJ**: NIH (R01HL140067); Hugh A. McAllister Research Foundation. **AMJ**: Intramural Research Program of the National Institute of Environmental Health Sciences, the National Institutes of Health [Z01-ES-101586].

## Materials and Methods

### Experimental animals

Heterozygous RORα^sg/sg^ mice on a C57BL/6J background were purchased from Jackson lab and maintained as previously described (Beak *et al.*, 2019). Homozygous RORα^sg/sg^ mice, the products of heterozygous breeding, and WT littermates were used in all experiments at 12-16 weeks of age. Homozygous cardiomyocyte-specific RORα knockout mice (RORα CMKO) were generated by crossing floxed RORα mice with LoxP sites flanking exon 9 (Anton Jetten, NIEHS, Durham, NC) with αMHC-Cre mice (Dale Abel, University of Iowa). All animal studies followed the NIH Guide for the Care and Use of Laboratory Animals and animal protocols were approved by the NIEHS Animal Care and Use Committee and the University of North Carolina Institutional Animal Care and Use Committee.

### Transmission Electron Microscopy (TEM)

Cardiac sections were observed with a LEO EM910 TEM operating at 80 kV (LEO Electron Microscopy, Thornwood, NY) and photographed with a Gatan Orius SC1000 CCD Digital Camera and Digital Micrograph 3.11.0 (Gatan, Pleasanton, CA). Mitochondrial cross-sectional area was measured using NIH ImageJ at magnifications of ×5000 to ×10,000. The global scale was set according to the image-specific scale generated by the Gatan camera output. An average of 2500 mitochondria were analyzed from 30 fields from multiple levels from three hearts per mouse cohort.

### Microarray analysis

Microarray analysis was performed to compare cardiac gene expression between WT (n=5) and RORα^sg/sg^ (n=4) mice. RNA was amplified using NuGEN Ovation Pico WTA System (NuGEN). Gene expression analysis was conducted using Agilent Whole Mouse Genome 4×44 multiplex format oligo arrays (Agilent Technologies 014868) following the Agilent 1-color microarray-based gene expression analysis protocol. Starting with 500ng of total RNA, Cy3 labeled cRNA was produced according to manufacturer’s protocol. For each sample, 1.65ug of Cy3 labeled cRNA was fragmented and hybridized for 17 hours in a rotating hybridization oven. Slides were washed and then scanned with an Agilent Scanner. Data was obtained using the Agilent Feature Extraction software (v9.5), using the 1-color defaults for all parameters. The Agilent Feature Extraction Software performed error modeling, adjusting for additive and multiplicative noise. The resulting data were processed using the Rosetta Resolver^®^ system version 7.2 (Rosetta Biosoftware, Kirkland, WA). In order to identify differentially expressed probes, analysis of variance (ANOVA) was used to determine if there was a statistical difference between the means of groups. In addition, we used a multiple test correction to reduce the number of false positives. Specifically, an error weighted ANOVA and Benjamini-Hochberg multiple test correction with a p value of p<0.01 was performed using Rosetta Resolver (www.rosettabio.com). The data presented in this publication have been deposited in the NCBI Gene Expression Omnibus at http://www.ncbi.nlm.nih.gov/geo/#GSE98892.

### Hypoxia chamber

Mice were housed in a hypoxia chamber (Oxycycler) with internal dimension (76×51×51 cm) sufficient to hold 4 cages with 4 mice per cage as previously described (Zhang *et al*, 2020). Inflow rate (nitrogen mixed with oxygen or room air) was ~3.1 ft^3^/hr and 4 holes (0.7 cm diameter) were opened at the bottom of each of the chamber’s 3 sides. Drierite (#21909-5000, Acros, Fair Lawn, NJ) and soda lime (#36596, Alfa Aesar, Haverhill, MA) were placed at the bottom of the chamber in 12×10×5 cm trays (~260g and 200g, respectively). CO_2_ was measured using a Cozir Wide-Range sensor and GasLab software, (#CM-0123 CO2meter.com, Ormond Beach, Fl). The hypoxia chambers have internal fans to prevent gradients in composition of the atmospheres. CO2 was automatically recorded at 5 min intervals. FIO_2_ was lowered from 21% to 10% (hypoxia) in ~3 h. Control (21% O_2_, normoxia) mice were maintained in an identical chamber. Hypoxia chambers were generously provided by Dr. James Faber.

### Histology and immunofluorescence microscopy

Mice were heparinized and the heart was perfused with 10 mL PBS followed by 20 mL of 4% PFA/PBS through a 23G butterfly needle, then excised and placed in 4% PFA/PBS for 24h prior to transfer to 70% EtOH. For immunohistochemistry, hearts were fixed overnight in 4% PFA/PBS, incubated in 30% sucrose/PBS, then frozen in O.T.C. medium (Tissue-Tek, Hatfield, PA). Frozen sections (10 microns), obtained with a Leica cryostat (Leica, Buffalo Grove, IL), were placed on glass slides, dried at room temperature, and then incubated with primary antibodies. After washing, the sections were incubated for 3 hours at room temperature with anti-mouse, anti-rabbit, anti-goat, or anti-rat Alexa Fluor-488, Fluor-594 or Fluor-647 conjugated secondary antibodies (1:1,000, Life Technologies, Grand Island, NY). Fluorescence was observed with a Zeiss LSM880 confocal microscope.

### Quantitative real-time reverse transcriptase PCR (qRT-PCR)

Total mouse heart RNA was isolated with an RNeasy mini kit (Qiagen, Valencia, CA) or RNAqueous^®^ Micro RNA isolation kit (Ambion, Austin, TX) following the manufacturer’s instructions. RNA was reverse-transcribed using a High-Capacity cDNA Archive Kit (Applied Biosystems, Foster City, CA). qRT-PCR reactions were carried out in triplicate in a LightCycler^®^ 480 System (Roche, Indianapolis, IN, USA) using either probe/primer sets or SYBR Green I (see Primer List). Relative quantitation of PCR products used the ΔΔCt method relative to two validated reference genes (Tbp1 and Polr2a). All probes and primers were from Roche or Thermofisher.

### Immunoblotting

Whole tissue or cell lysates were produced in RIPA buffer supplemented with PhosSTOP (Roche Diagnostics Corporation, Indianapolis, IN, USA) and protease inhibitor cocktail (Roche Diagnostics Corporation). Subsequently samples were incubated in 4× LDS sample buffer with 2% β-mercaptoethanol, for 10 min at 70 °C. SDS–PAGE and immunoblotting were performed using the 4-12% NuPage gel system (Life Technologies, Foster City, CA, USA). Membranes were blocked in 5% milk/TBS-Tween, incubated in primary antibody overnight at 4°C, then secondary HRP-conjugated antibodies for 1h at room temperature. Images were generated using Amersham ECL Select Western Blotting Detection Reagent (GE Healthcare life sciences, Marlborough, MA, USA) and the MultiDoc-It™ Imaging System (UVP gel image system, Upland, CA, USA).

### Antibodies

RORα (GTX100029, 1:1000), SQSTM1/p62 (GTX629890, 1:1000), PINK1 (GTX107851, 1:1000), LC3B (GTX127375, 1:1000), Ambra1 (GTX55507, 1:1000), and HIF1α (GTX628480) were from Genetex (Irvine, CA, USA). VDAC1 (#4661, 1:000), HA (#3724, 1:1000), LC3B (#83506, 1:500), and Hsp60 (#12165, 1:1000) were from Cell Signaling Technology (Danvers, MA, USA). GAPDH (MAB374 clone 6C5, 1: 10,000, Millipore), Caveolin-3 (sc-5310, 1:1000), Parkin (sc-32282, 1:1000), and Bnip3 (sc-56167, 1:1000) were from Santa Cruz Biotechnology (Dallas, TX, USA). Polyclonal goat anti-rabbit IgG–horseradish-peroxidase (HRP; A9169, 1:5,000), polyclonal rabbit anti-mouse IgG–HRP (A9044, 1:5,000), polyclonal rabbit anti-goat IgG-HRP (A5420, 1:5,000) were from Sigma Aldrich (St. Louis, MO, USA).

### Neonatal rat ventricular myocyte (NRVM) cultures, immunocytochemistry and lentiviral infections

Female Sprague-Dawley rats and newborn litters were from Charles River Laboratories (Wilmington, MA, USA). NRVMs were isolated as previously described (Simpson, 1983). Experiments were carried out after 36-48 hours of serum starvation in the presence of insulin, transferrin, and BrdU. NRVM hypoxia was induced by incubation in a 37°C hypoxia chamber with 1% O_2_ after infection with lentivirus containing either scrambled short hairpin RNA (shControl) or shRNA specifically targeting rat RORα (iO51217 or iV051217, abm, Richmond, BC, Canada).

shRORα target sequences: TGTCATTACGTGTGAAGGCTGCAAGGGCT, ACCTACAACATCTCAGCCAATGGGCTGAC, GGACTGGACATCAATGGGATCAAACCCGA, AGAGGTGATGTGGCAGTTGTGTGCTATCA

### Mitochondrial fractionation

Mitochondria were isolated from frozen heart tissue using a Mitochondria isolation kit (ab110168, abcam). Tissue was briefly washed in isolation buffer, dried with Whatman filter paper, weighed, placed in a glass beaker, thoroughly minced then homogenized with a Dounce homogenizer. The homogenate was centrifuged at 1,000g for 10 min at 4°C. The supernatant was centrifuged at 12,000g for 15 min at 4°C then saved as crude cytosolic and nuclear fractions and the pellet was centrifuged again at 12,000g for 15min after resuspending in isolation buffer and protease inhibitor for further mitochondrial purification. The homogenized pellets were re-suspended in isolation buffer with protease and phosphatase inhibitor cocktails (Roche, Indianapolis, IN). The protein concentration was measured by BCA protein assay kit (Thermo Scientific, Waltham, MA).

### ATP bioluminescence assay

ATP content was determined using a luciferin-luciferase ATP assay (ThermoFisher Scientific A22066) according to the manufacturer’s protocol. All reagents were placed in ice before the experiments were carried out. The ATP solution (100 nM) was prepared by mixing 50μl of a stock solution (5 mM) with 950 μl of deionized water (DW), and buffer in a microcentrifuge tube. The mixture was then transferred to a 96-well microplate for measurement of ATP bioluminescence. In brief, 100 μl of the ATP solution was mixed with 100 μl of the luciferin–luciferase reagent (10 mg/ml; reconstituted with DW) in each well of the microplate. ATP bioluminescence was measured immediately using a microplate luminometer. The integration time of the luminometer was set at 1 s with normal gain.

### Mitochondrial membrane potential

Mitochondrial membrane potential in NRVMs was determined by 5, 5′, 6, 6′-tetrachloro-1, 1′, 3, 3′-tetraethylbenzimidazolylcarbocyanine iodide (JC-1) reduction. Cells were incubated with JC-1 (Cambridge, MA, Abcam) according to the manufacturer's protocol. In brief, serum-starved NRVMs were incubated in the normoxia chamber with 21% O**2**or in the hypoxia chamber with 1% after infection with shControl or shRORα. JC-1 2 μM was added to each well for 30 min. Cells were washed once with medium then analyzed by plate reader (CLARIOstar, BMG LABTECH, Germany). JC-1 green fluorescence was excited at 488 nm and emission was detected using a 530 ± 40 nm filter. JC-1 red fluorescence was excited at 488 nm and emission was detected using a 613 ± 20 nm filter.

### Succinate dehydrogenase (SDH) activity assay

SDH activity was measured using an SDH Activity Colorimetric Assay Kit (MBS2540432) purchased from MyBiosource Co. (San Diego, CA). After correcting for the background, SDH activity in NRVM lysates was determined.

### LDH release assay

For measurement of cytotoxicity, NRVMs were seeded onto 96-well plates. After treatment, 50 μl of culture medium from each well was collected and transferred to another 96-well culture plate. 50 μl of LDH reagent (Thermo Fisher scientific, C20300) was then added to each well and the solution incubated at room temperature for 30 min. After the reaction was complete, absorbance at 490 nm was measured by an ELISA reader.

### Citrate synthase activity

Citrate synthase activity assay kit (MAK 193, Millipore sigma, St. Louis, MO) was used in NRVMs incubated at 21% or 1% O2 after infection with shControl- or shRORα-lentivirus.

### Mitochondrial ROS measurement

The production of superoxide by the mitochondria of NRVMs incubated at 21% or 1% O_2_ after infection with shControl- or shRORα-lentivirus was measured using MitoSOX™ Red mitochondrial superoxide indicator (M36008, Thermo Fisher scientific).

### Oxygen consumption rate (OCR) assay

OCR was measured using an extracellular oxygen consumption assay kit (ab197243; Abcam, Cambridge, UK) following the manufacturer’s protocol. NRVMs in a 96-well black plate were incubated at 21% or 1% O_2_ after infection with shControl- or shRORα-lentivirus in triplicates for each condition. A negative control was prepared with reaction buffer without cells. All wells were covered with two drops of pre-heated mineral oil to block the back diffusion of oxygen. OCR was calculated based on fluorescent signals (excitation of 380 nm/emission of 650 nm), measured in 1.5 min intervals for 1.5 h using a plate reader (CLARIOstar, BMG LABTECH, Germany). The experiment was repeated two times independently.

### Keima with Mitochondrial Localization Signal

Keima with mitochondrial localization signal (Mito-Keima), a mitochondrially localized pH-indicator protein (Katayama *et al*, 2011), was generously supplied by Junichi Sadoshima (Rutgers New Jersey Medical School). The method used to detect lysosomal delivery of Mito-Keima has been described (Shirakabe *et al.*, 2016). Briefly, NRVMs were infected with shControl- or shRORα-lentivirus for 24 hr then infected with mtKeima-virus for visualizing mitophagy (543 nm and 458 nm) under fluorescent microscopy. Relative fluorescent intensity (ratio of 543nm/458nm) was determined by Image J.

### Treatment with RORα inverse agonist (SR3335) and agonist (SR1078)

NRVMs were treated with vehicle or RORα inverse agonist, SR3335 20μM (Kumar *et al*, 2011) or agonist, SR1078 2μM(Wang *et al.*, 2016) for 6 hr and then incubated in a hypoxia chamber (1% O_2_) after pretreatment with BFA (100nM) for 2 hr.

### Chromatin immunoprecipitation-PCR

To demonstrate direct transcriptional regulation of Caveolin-3 by RORα, adult cardiomyocytes isolated from 3-4 month old mice were washed in PBS containing protease and phosphatase inhibitor cocktail (PIC; Thermo Scientific) and then fixed in 1% formaldehyde at 25°C for 8 min for cross-linking and followed by quenching by 0.125 M glycine at 25°C for 8 min as previously described. (Beak *et al.*, 2019) The amount of chromatin immunoprecipitated DNA (ChIPed-DNA) relative to each input DNA was determined by quantitative PCR in triplicate with primers amplifying consensus RORE (AGGTCA) or a nontargeting sequence in the mouse Caveolin-3 promoter.

Chromatin immunoprecipitation (ChIP)-PCR primers were:

Caveolin-3-4.2k, forward 5’-GCCAAGAGGGATCAGCAATA-3’ and reverse 5’-CAAGGGAAGGCATTTGACAT-3’;

Caveolin-3-0.5k, forward 5’-CCCCATGGCCTTAGTTTGAT-3’ and reverse 5’-TTATTTGCGTGCAAGTGAGC-3’;

Caveolin-3-0.1k, forward 5’-CAGGGTGGGAAGACTCTTGA-3’ and reverse 5’-GTCCCAGCTCTGTCTCTTGC-3’.

### Generation of doxycycline-inducible H9c2 cell line and immunocytochemistry

An H9c2 cell line with doxycycline-inducible RORα4 was adapted from a published report. (Jeon *et al*, 2019) Briefly, RORα4 tagged with N-terminus FLAG and C-terminus HA (FLAG-RORα4-HA) was inserted into lentiviral vector pIND20, and viral soup was generated in HEK293T cells. H9c2 cells were infected with lentiviral pIND20-FLAG-RORα4-HA and stable cells were selected by 500 μg/ml of G418. Induction of RORα4 was examined by Western blot using HA antibody after treatment of doxycycline. For the Cav-3 rescue experiment, we infected H9c2s with shRNA against RORα or scrambled shRNA for 48hrs then transfected with an EGFP-Cav-3 plasmid (Addgene plasmid #68396) or empty EGFP plasmid and an mRaspberry-Mito-7 plasmid (Addgene #55931) to localize mitochondria. For visualization of mitophagy, H2c2s were stained with LC3B and LysoTracker (Thermo Fisher Scientific, L7525).

### Statistics

All results are presented as mean ± SEM. Comparisons were made in GraphPad Prism (San Diego, CA) using unpaired t-test (2 groups) or two-way ANOVA (Type III sum of squares) with Tukey’s post-hoc analysis (4 groups).

---

**Supplemental Figure S1.**
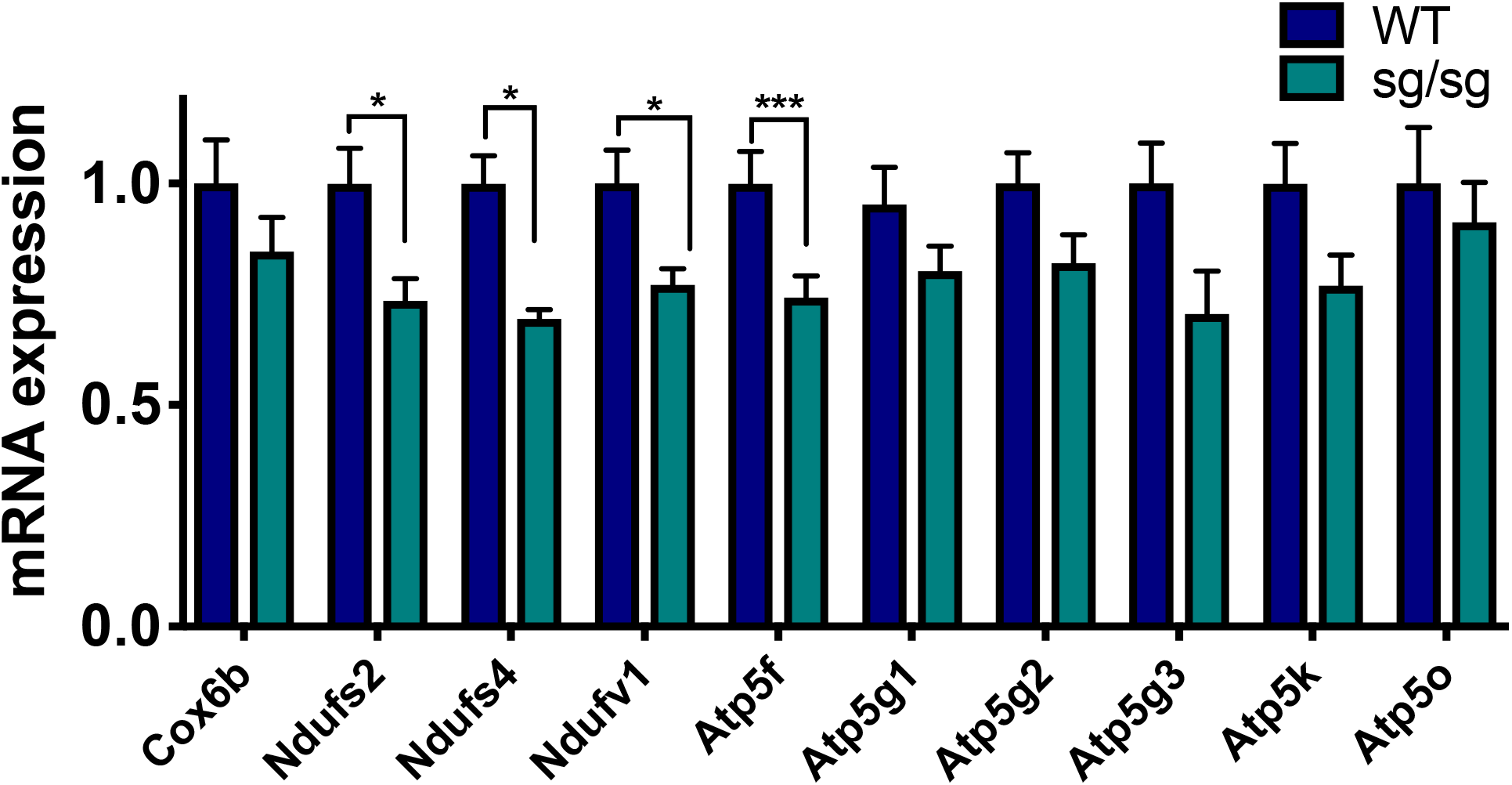
Quantitative reverse transcriptase PCR compared abundance of selected electron transport chain transcripts in wild type (WT) and RORα^sg/sg^ (sg/sg) mice (n=6 per group).

**Supplemental Figure S2.**
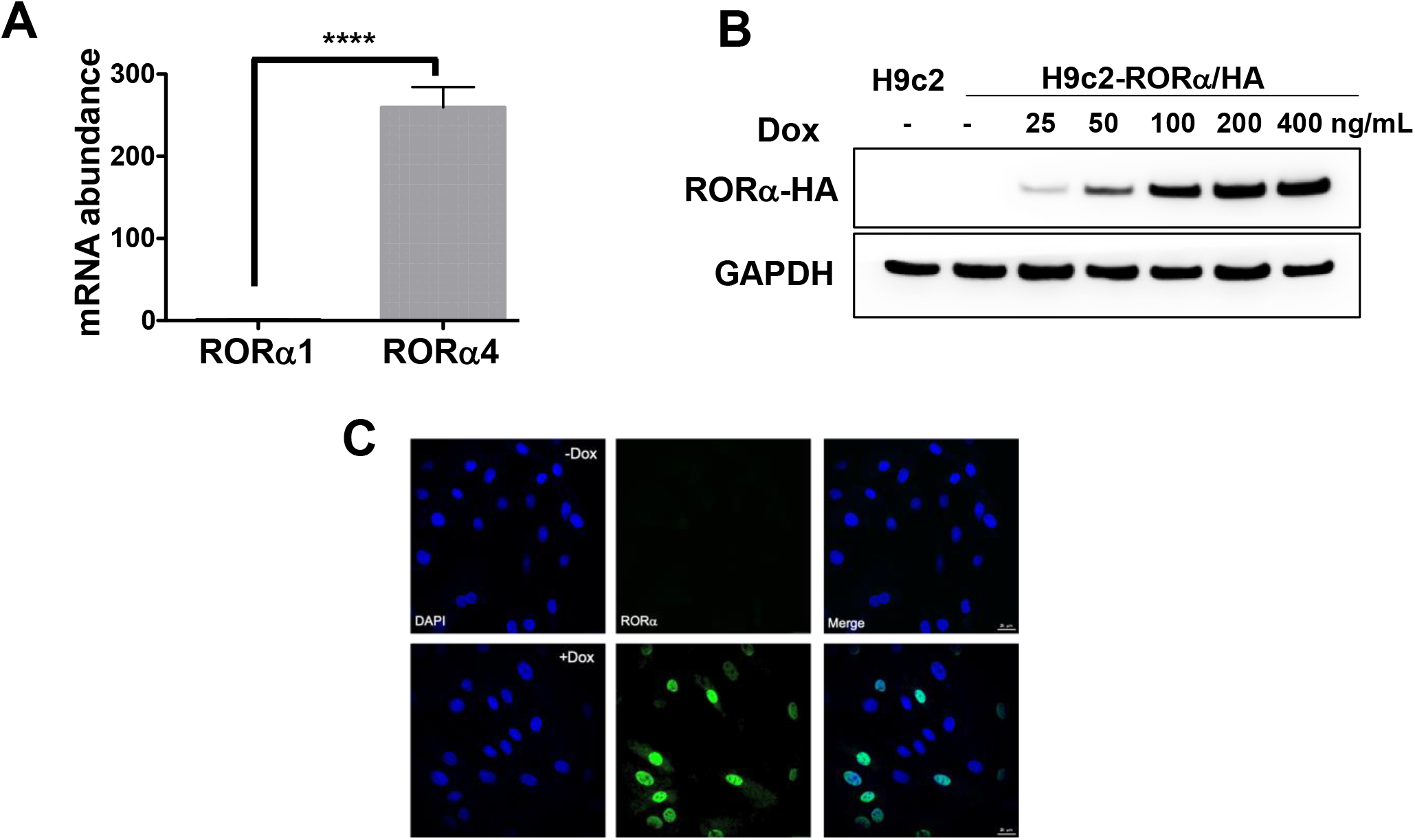
**(A)** Quantitative reverse transcriptase PCR for RORα subtypes in wild type mouse heart (n = 5) **(B)** Immunoblotting of cell lysates from H9c2 and H9c2-Flag/RORα4/HA with varying concentrations of doxycycline. **(C)** Immunofluorescence staining of HA antibody (RORα4, green) and DAPI (blue) in H9c2-Flag/RORα4/HA with or without doxycycline.

## REFERENCES

Beak JY, Kang HS, Huang W, Myers PH, Bowles DE, Jetten AM, Jensen BC (2019) The nuclear receptor RORalpha protects against angiotensin II-induced cardiac hypertrophy and heart failure. Am J Physiol Heart Circ Physiol 316: H186–H200

Cadenas S, Aragones J, Landazuri MO (2010) Mitochondrial reprogramming through cardiac oxygen sensors in ischaemic heart disease. Cardiovasc Res 88: 219–228

Chauvet C, Bois-Joyeux B, Berra E, Pouyssegur J, Danan JL (2004) The gene encoding human retinoic acid-receptor-related orphan receptor alpha is a target for hypoxia-inducible factor 1. Biochem J 384: 79–85

Cook DN, Kang HS, Jetten AM (2015) Retinoic Acid-Related Orphan Receptors (RORs): Regulatory Functions in Immunity, Development, Circadian Rhythm, and Metabolism. Nucl Receptor Res 2

Fridolfsson HN, Kawaraguchi Y, Ali SS, Panneerselvam M, Niesman IR, Finley JC, Kellerhals SE, Migita MY, Okada H, Moreno AL et al. (2012) Mitochondria-localized caveolin in adaptation to cellular stress and injury. FASEB J 26: 4637–4649

He B, Zhao Y, Xu L, Gao L, Su Y, Lin N, Pu J (2016) The nuclear melatonin receptor RORalpha is a novel endogenous defender against myocardial ischemia/reperfusion injury. J Pineal Res 60: 313–326

Horikawa YT, Panneerselvam M, Kawaraguchi Y, Tsutsumi YM, Ali SS, Balijepalli RC, Murray F, Head BP, Niesman IR, Rieg T et al. (2011) Cardiac-specific overexpression of caveolin-3 attenuates cardiac hypertrophy and increases natriuretic peptide expression and signaling. J Am Coll Cardiol 57: 2273–2283

Horikawa YT, Patel HH, Tsutsumi YM, Jennings MM, Kidd MW, Hagiwara Y, Ishikawa Y, Insel PA, Roth DM (2008) Caveolin-3 expression and caveolae are required for isoflurane-induced cardiac protection from hypoxia and ischemia/reperfusion injury. J Mol Cell Cardiol 44: 123–130

Huang da W, Sherman BT, Lempicki RA (2009a) Bioinformatics enrichment tools: paths toward the comprehensive functional analysis of large gene lists. Nucleic Acids Res 37: 1–13

Huang da W, Sherman BT, Lempicki RA (2009b) Systematic and integrative analysis of large gene lists using DAVID bioinformatics resources. Nat Protoc 4: 44–57

Jeon K, Kumar D, Conway AE, Park K, Jothi R, Jetten AM (2019) GLIS3 Transcriptionally Activates WNT Genes to Promote Differentiation of Human Embryonic Stem Cells into Posterior Neural Progenitors. Stem Cells 37: 202–215

Kassan A, Pham U, Nguyen Q, Reichelt ME, Cho E, Patel PM, Roth DM, Head BP, Patel HH (2016) Caveolin-3 plays a critical role in autophagy after ischemia-reperfusion. Am J Physiol Cell Physiol 311: C854–C865

Katayama H, Kogure T, Mizushima N, Yoshimori T, Miyawaki A (2011) A sensitive and quantitative technique for detecting autophagic events based on lysosomal delivery. Chem Biol 18: 1042–1052

Kim EJ, Yoo YG, Yang WK, Lim YS, Na TY, Lee IK, Lee MO (2008) Transcriptional activation of HIF-1 by RORalpha and its role in hypoxia signaling. Arterioscler Thromb Vasc Biol 28: 1796–1802

Kim HJ, Han YH, Na H, Kim JY, Kim T, Kim HJ, Shin C, Lee JW, Lee MO (2017) Liver-specific deletion of RORalpha aggravates diet-induced nonalcoholic steatohepatitis by inducing mitochondrial dysfunction. Sci Rep 7: 16041

Kong CHT, Bryant SM, Watson JJ, Roth DM, Patel HH, Cannell MB, James AF, Orchard CH (2019) Cardiac-specific overexpression of caveolin-3 preserves t-tubular ICa during heart failure in mice. Exp Physiol 104: 654–666

Kubli DA, Gustafsson AB (2012) Mitochondria and mitophagy: the yin and yang of cell death control. Circulation research 111: 1208–1221

Kubli DA, Zhang X, Lee Y, Hanna RA, Quinsay MN, Nguyen CK, Jimenez R, Petrosyan S, Murphy AN, Gustafsson AB (2013) Parkin protein deficiency exacerbates cardiac injury and reduces survival following myocardial infarction. J Biol Chem 288: 915–926

Kumar N, Kojetin DJ, Solt LA, Kumar KG, Nuhant P, Duckett DR, Cameron MD, Butler AA, Roush WR, Griffin PR et al. (2011) Identification of SR3335 (ML-176): a synthetic RORalpha selective inverse agonist. ACS Chem Biol 6: 218–222

Lau P, Fitzsimmons RL, Raichur S, Wang SC, Lechtken A, Muscat GE (2008) The orphan nuclear receptor, RORalpha, regulates gene expression that controls lipid metabolism: staggerer (SG/SG) mice are resistant to diet-induced obesity. J Biol Chem 283: 18411–18421

Lau P, Nixon SJ, Parton RG, Muscat GE (2004) RORalpha regulates the expression of genes involved in lipid homeostasis in skeletal muscle cells: caveolin-3 and CPT-1 are direct targets of ROR. J Biol Chem 279: 36828–36840

Mauvezin C, Neufeld TP (2015) Bafilomycin A1 disrupts autophagic flux by inhibiting both V-ATPase-dependent acidification and Ca-P60A/SERCA-dependent autophagosome-lysosome fusion. Autophagy 11: 1437–1438

Moyzis AG, Sadoshima J, Gustafsson AB (2015) Mending a broken heart: the role of mitophagy in cardioprotection. Am J Physiol Heart Circ Physiol 308: H183–192

Nah J, Yoo SM, Jung S, Jeong EI, Park M, Kaang BK, Jung YK (2017) Phosphorylated CAV1 activates autophagy through an interaction with BECN1 under oxidative stress. Cell Death Dis 8: e2822

Ratajczak P, Damy T, Heymes C, Oliviero P, Marotte F, Robidel E, Sercombe R, Boczkowski J, Rappaport L, Samuel JL (2003) Caveolin-1 and −3 dissociations from caveolae to cytosol in the heart during aging and after myocardial infarction in rat. Cardiovasc Res 57: 358–369

Sharifi MN, Mowers EE, Drake LE, Macleod KF (2015) Measuring autophagy in stressed cells. Methods Mol Biol 1292: 129–150

Shirakabe A, Fritzky L, Saito T, Zhai P, Miyamoto S, Gustafsson AB, Kitsis RN, Sadoshima J (2016) Evaluating mitochondrial autophagy in the mouse heart. J Mol Cell Cardiol 92: 134–139

Simpson P (1983) Norepinephrine-stimulated hypertrophy of cultured rat myocardial cells is an alpha_1_-adrenergic response. J Clin Invest 72: 732–738

Tang Z, Scherer PE, Okamoto T, Song K, Chu C, Kohtz DS, Nishimoto I, Lodish HF, Lisanti MP (1996) Molecular cloning of caveolin-3, a novel member of the caveolin gene family expressed predominantly in muscle. J Biol Chem 271: 2255–2261

Tsutsumi YM, Horikawa YT, Jennings MM, Kidd MW, Niesman IR, Yokoyama U, Head BP, Hagiwara Y, Ishikawa Y, Miyanohara A et al. (2008) Cardiac-specific overexpression of caveolin-3 induces endogenous cardiac protection by mimicking ischemic preconditioning. Circulation 118: 1979–1988

Vega RB, Kelly DP (2017) Cardiac nuclear receptors: architects of mitochondrial structure and function. J Clin Invest 127: 1155–1164

Wang Y, Billon C, Walker JK, Burris TP (2016) Therapeutic Effect of a Synthetic RORalpha/gamma Agonist in an Animal Model of Autism. ACS Chem Neurosci 7: 143–148

Zhang H, Rzechorzek W, Aghajanian A, Faber JE (2020) Hypoxia induces de novo formation of cerebral collaterals and lessens the severity of ischemic stroke. J Cereb Blood Flow Metab 40: 1806–1822

Zhao Y, Xu L, Ding S, Lin N, Ji Q, Gao L, Su Y, He B, Pu J (2017) Novel protective role of the circadian nuclear receptor retinoic acid-related orphan receptor-alpha in diabetic cardiomyopathy. J Pineal Res 62

Zhou B, Tian R (2018) Mitochondrial dysfunction in pathophysiology of heart failure. J Clin Invest 128: 3716–3726

Zhou Q, Peng X, Liu X, Chen L, Xiong Q, Shen Y, Xie J, Xu Z, Huang L, Hu J et al. (2018) FAT10 attenuates hypoxia-induced cardiomyocyte apoptosis by stabilizing caveolin-3. J Mol Cell Cardiol 116: 115–124

